# Transcriptional and translational responsiveness of the *Neisseria gonorrhoeae* type IV secretion system to conditions of host infections

**DOI:** 10.1101/2021.02.12.431057

**Authors:** Melanie M. Callaghan, Amy K. Klimowicz, Abigail C. Shockey, John Kane, Caitlin S. Pepperell, Joseph P. Dillard

## Abstract

The type IV secretion system of *Neisseria gonorrhoeae* translocates single-stranded DNA into the extracellular space, facilitating horizontal gene transfer and initiating biofilm formation. Expression of this system has been observed to be low under laboratory conditions, and multiple levels of regulation have been identified. We used a translational fusion of *lacZ* to *traD*, the gene for the type IV secretion system coupling protein, to screen for increased type IV secretion system expression. We identified several physiologically relevant conditions, including surface adherence, decreased manganese or iron, and increased zinc or copper, which increase gonococcal type IV secretion system protein levels through transcriptional and/or translational mechanisms. These metal treatments are reminiscent of the conditions in the macrophage phagosome. The ferric uptake regulator, Fur, was found to repress *traD* transcript levels, but to also have a second role, acting to allow TraD protein levels to increase only in the absence of iron. To better understand type IV secretion system regulation during infection, we examined transcriptomic data from active urethral infection samples from five men. These data demonstrated differential expression of 20 of 21 type IV secretion system genes during infection, indicating upregulation of genes necessary for DNA secretion during host infection.

## Introduction

*Neisseria gonorrhoeae*, the bacterial agent of the sexually transmitted infection gonorrhea, establishes thriving infections on human mucosal surfaces (e.g., the urogenital tract, rectum, nasopharynx, and eyes). Symptomatic gonorrhea infections are characterized by robust inflammatory responses, with extensive influx of both activated neutrophils and macrophages (Sintsova et al. 2014; Château and Seifert 2016). *N. gonorrhoeae* have been shown to invade and survive in epithelial cells and neutrophils and to cause programmed cell death in macrophages (King et al. 1978; Apicella et al. 1996; Merz et al. 1996; Harvey et al. 2001; Criss et al. 2009; Château and Seifert 2016; Ritter and Genco 2018). The wide variety of tissues and cell types that *N. gonorrhoeae* interacts with throughout the course of any given infection reflects the bacterium’s capacity to respond to diverse environments.

Many *N. gonorrhoeae* strains possess the 59kb gonococcal genetic island (GGI) on their ∼2.2Mb chromosome (Dillard and Seifert 2001). This island has been found in 60-80% of isolates and encodes a unique type IV secretion system (T4SS) (Dillard and Seifert 2001; Hamilton et al. 2005; Callaghan et al. 2017; Shockey 2019). T4SSs are a diverse subset of protein nanomachines that can be used for protein secretion, conjugation, DNA uptake, and in the case of *N. gonorrhoeae*, DNA secretion (Backert and Grohmann 2017). The gonococcal T4SS exports single-stranded DNA directly into the extracellular space independent of contact with a host or neighboring cell (Dillard and Seifert 2001; Hamilton et al. 2005; Salgado-Pabón et al. 2007). This unique method of DNA secretion has been shown to serve many purposes in *N. gonorrhoeae* populations. The T4SS plays a role in biofilm establishment, largely by contributing an abundance of “sticky” DNA to nucleate and contribute to the biofilm matrix (Zweig et al. 2014). The secreted DNA has also been observed to be taken up by neighboring gonococci and stably incorporated into the chromosome, acting as a mechanism of horizontal gene transfer between gonococcal populations (Hamilton et al. 2001; Hamilton and Dillard 2006; Kohler et al. 2013). This process has been implicated in the spread of antibiotic resistance in gonococcal populations; GGI-containing strains of gonococci are more likely to have increased resistance to antibiotics (Harrison et al. 2016). While detectable genetic exchange does occur under laboratory conditions, detecting T4SS proteins and secreted DNA substrates in the laboratory has proven challenging due to low levels of expression in this environment (Dillard and Seifert 2001; Hamilton et al. 2005; Kohler et al. 2013; Ramsey et al. 2014; Ramsey et al. 2015).

Only 21 of the 66 GGI genes (located on only four of twelve identified operons) are necessary for the T4SS to be active (Figure 1A) (Hamilton et al. 2005; Jain et al. 2012; Pachulec et al. 2014; Remmele et al. 2014). Many of the gonococcal T4SS proteins, especially the structural components, are homologous to those of F-like plasmid-encoded conjugation systems (Pachulec et al. 2014). The remainder of the GGI is largely comprised of genes of unknown function (Hamilton et al. 2005; Pachulec et al. 2014). While some regulatory components have been characterized within the GGI, no homologues of F-like T4SS regulators have been found and a holistic picture of regulation is lacking in this system (Salgado-Pabón et al. 2010; Ramsey et al. 2014; Ramsey et al. 2015). The aforementioned low levels of T4SS expression have compounded the challenge of investigating regulation of the T4SS.

**Figure 1.**
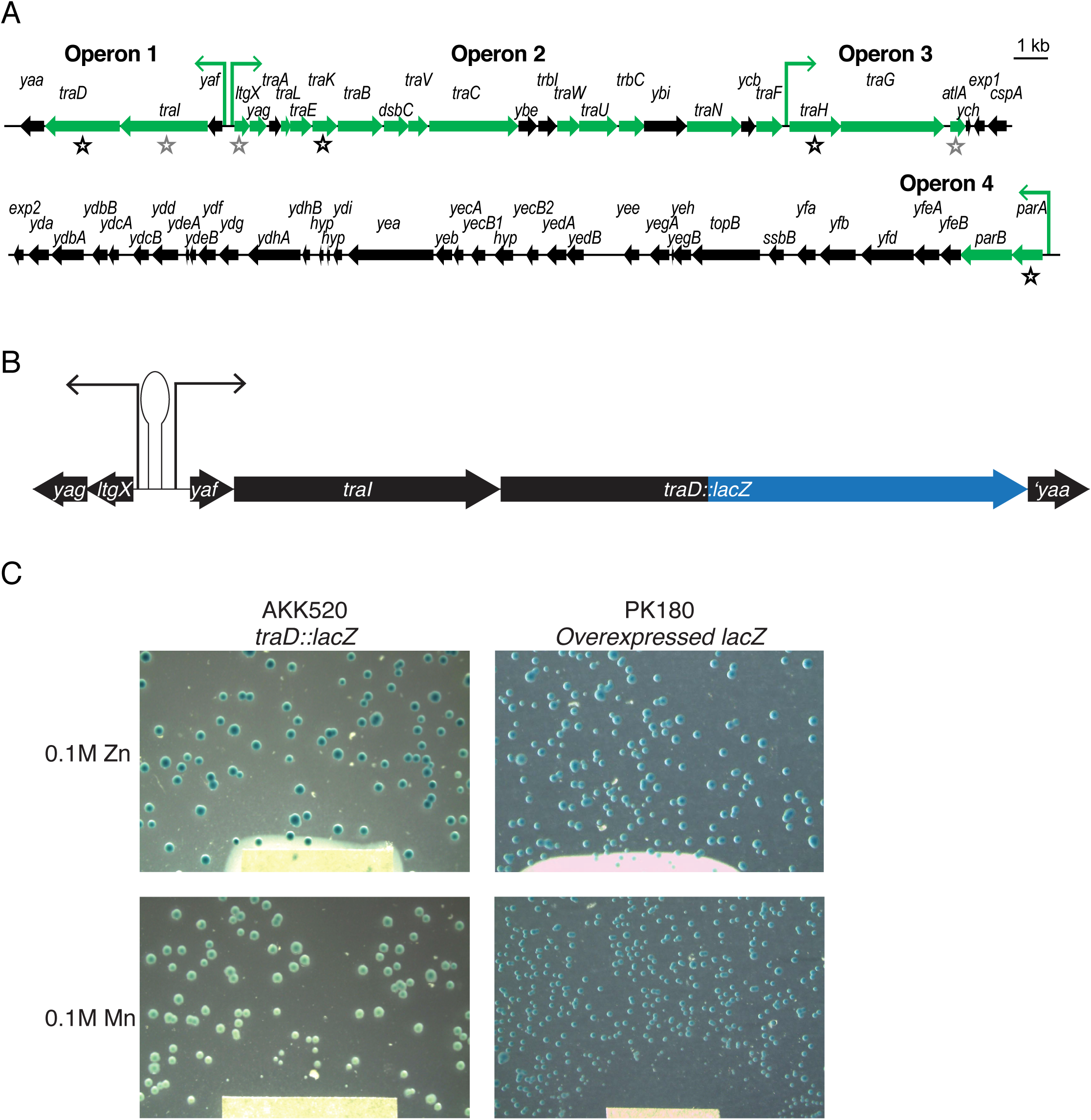
The GGI and the *traD::lacZ* reporter construct. (A) Map of the gonococcal genetic island (GGI). Genes necessary for secretion are shown in green. Transcripts measured in qRT-PCR experiments are denoted with stars. Promoters (green arrows) are marked for operons necessary for secretion only. (B) Gene map of *traD::lacZ* translational fusion reporter construct. (C) Representative images of *N. gonorrhoeae* colonies expressing LacZ in the presence of zinc or manganese-soaked disks. The left column shows TraD-LacZ expression from the native *traD* locus. Note color differences between colonies directly proximal to suffused disk compared to those more distant from disks. The right column shows colonies with LacZ overexpressed from a complementation site, which is neither zinc nor manganese responsive.

In this study, we find that the low expression seen in the laboratory is not representative of T4SS expression during host infection. We hypothesize that *N. gonorrhoeae* may invest in building the T4SS DNA pump only in response to signals specific to host environmental niches. Similar temporospatial determinants of expression have been identified in other T4SSs. For example, intracellular growth conditions signal *Legionella* to express a ubiquitin ligase that regulates T4SS protein levels via the host cell proteasome, and *Bartonella henselae*’s VirB/D4 T4SS is induced by interwoven signals from host cell pH and bacterial stringent response (Kubori et al. 2010; Québatte et al. 2013). Building on these observations, we aimed to determine how the environment of infection affects T4SS expression in GGI^+^ gonococci.

By screening an array of compounds, we identified several conditions that alter expression of the type IV coupling protein, TraD. We followed up this screen with quantitative transcript and protein measurements, and determined that iron chelation, zinc, and copper all enhance TraD expression, whereas manganese downregulates T4SS expression. We also found that surface adherence upregulates gene expression across the GGI. Our findings, along with the previous reports from our laboratory that piliation enhances type IV secretion (T4S), (Salgado-Pabón et al. 2010), are consistent with a model of increased GGI expression in response to specific conditions encountered within the human host. We tested this hypothesis by performing a transcriptomic analysis using publicly available data from active urethral infections (Nudel et al. 2018) and discovered significant changes in GGI expression during infection, including increased expression of multiple T4SS genes.

## Results

### Screen for conditions affecting TraD expression

Gonococci isolated from infections are always piliated, and piliated gonococci are more sensitive to antibiotics in the growth medium than non-piliated variants (Swanson et al. 1987; Gibbs et al. 1989). Moreover, previous studies have demonstrated that piliated gonococcal strains secrete more DNA and exhibit increased expression of *traD*, encoding the type IV coupling protein ATPase; *traI*, encoding the type IV secretion relaxase; and the GGI-encoded hypothetical protein *ydhB* (Salgado-Pabón et al. 2010). Due to its increased expression in the more robustly secreting piliated cells and its necessity for secretion, we selected TraD as a representative T4SS protein and made a *traD::lacZ* translational fusion expressed at the native *traD* locus in gonococci (AKK520) (Salgado-Pabón et al. 2010; Pachulec et al. 2014) (Figure 1B). Using this LacZ reporter in piliated cells, we tested a panel of infection-relevant compounds on TraD expression using disk diffusion assays on agar plates containing X-gal (Table 1, Figure 1C).

**Table 1.** *N. gonorrhoeae* expressing the *traD::lacZ* reporter at the native *traD* locus were spread onto GCB+X-gal plates, and compounds of interest screened by disk diffusion assay with blue colony color employed as the metric of protein expression. “No effect” indicates no visible changes in colony color in response to the compound-suffused disk. ↑ indicates increased color with proximity to disk (or increased color in the tested mutant). ↓ indicates decreased color with proximity to disk. *Side-by-side comparison of mutant and wild-type bacteria, as opposed to disk diffusion, used to determine difference in blue color.

| Category | Test Condition | Description | Observed blue color |
| --- | --- | --- | --- |
| Carbon sources | Glucose | 40mg/mL | No effect |
|  | Lactate | 60% (w/w) sodium DL-lactate solution | No effect |
|  | Acetate | 3M NaOAc | No effect |
| Antibiotics | Ciprofloxacin | 8mg/mL | No effect |
|  | Streptomycin | 100mg/mL | No effect |
| Extracellular material | Spent medium | 10μL filter-sterilized supernatant, *N. gonorrhoeae* MS11 in cGCBL | No effect |
| DNA | DNA | *N. gonorrhoeae* gDNA | No effect |
|  | *recA4* * | *recA* loss of function allele (Seifert 1997) | ↑ |
| ROS/RNS | Hydrogen peroxide | 3% solution | ↑ |
|  | Nitrite | 500mM NaNO<sub>2</sub> | No effect |
|  | SNAP | NO donor, S-Nitroso-N-acetyl-DL-penicillamine, 90mM | No effect |
|  | GSNO | NO donor, S-Nitrosoglutathione, 50mM | No effect |
| Infection/transmission | Spermidine | 100mg/mL | No effect |
|  | Seminal plasma | ≥3 pooled donors per sample | ↑ |
| Metals | Iron | 1.2μM Fe(NO<sub>3</sub>)<sub>3</sub> | No effect |
|  | Desferal (Desferoxamine mesylate) | Iron chelator, 25mM | ↑ |
|  | *fur-1\** | *fur* point mutation, partially functional Fur | ↑ |
|  | Magnesium | 100mM MgCl<sub>2</sub> | No effect |
|  | Manganese | 100mM MnCl<sub>2</sub> | ↓ |
|  | Copper | 100mM CuSO<sub>4</sub> | ↑ |
|  | Cobalt | 100mM CoCl<sub>2</sub> | No effect |
|  | Zinc | 100mM ZnSO<sub>4</sub> | ↑ |
|  | EDTA | 250mM | ↑ |

Gonococci infect a variety of environmentally distinct niches, and it is possible that the T4SS is only active at certain infection sites. The host typically sequesters metal at sites of infection; pathogen killing via nutritional immunity strategies has been documented for iron, zinc, and manganese (Kehl-Fie and Skaar 2010; Damo et al. 2013; Zygiel et al. 2019). *N. gonorrhoeae* utilizes several TonB-dependent and ABC transport systems to obtain necessary metals, and modulates expression of these scavenging and import systems in response to metals in its environment (Cornelissen 2018; Maurakis et al. 2019). Supplemental iron had no visible effect on TraD (Table 1), however GCB agar has high iron content due to the addition of Kellogg’s supplements, so additional iron may have been superfluous for regulation. By contrast, the iron chelator desferal increased TraD reporter signal, suggesting that low iron availability increases TraD expression. EDTA increased TraD similarly (Table 1). We expanded our screen to include other metals. Solutions of copper or zinc each increased β-galactosidase activity in our screen (Table 1, Figure 1C). The only compound tested that visibly decreased signal was manganese (Table 1, Figure 1C). No effect on β-galactosidase activity was seen in a control strain constitutively expressing *lacZ* (Figure 1C). Based on these observations, we hypothesize that T4SS expression changes in response to stimuli including metal availability.

Next, we tested carbon sources metabolized by *N. gonorrhoeae*. Neither solutions of glucose, its preferred carbon source, nor lactate, an additional carbon source utilized for growth during infection (Morse and Williams 1979; Exley et al. 2007) altered TraD levels in the disk diffusion LacZ reporter assay (Table 1). Acetate, a metabolizable byproduct of growth (Hebeler and Morse 1976), also showed no effect on TraD levels (Table 1).

Transcriptional changes in response to sub-inhibitory levels of antibiotics have been identified in a number of bacterial pathogens, including *Salmonella enterica* serovar Typhimurium and *N. gonorrhoeae* (Goh et al. 2002; Arvidson et al. 2006). Clinical isolates of *N. gonorrhoeae* have demonstrated resistance to all classes of antibiotics, making antibiotic response an especially salient question for gene regulation (Unemo and Shafer 2014). However, we found no effect from ciprofloxacin or streptomycin on TraD expression in the disk diffusion assay (Table 1).

We also tested genomic DNA, the substrate of the T4SS, and spent medium from cultures of wild type GGI^+^ gonococci, which contains DNA from both autolysis and secretion as well as other growth and lysis products. Neither affected TraD expression (Table 1). Transformation of DNA, which may be facilitated by the T4SS, is speculated to provide material for DNA repair (Davidsen et al. 2004). We probed the role of repair in regulating T4S by testing a DNA recombination deficient mutant (*recA4,* (Seifert 1997)) in our disk diffusion assay. The recombination mutant did have stronger blue color compared to wild type *recA*^+^ cells, indicating some potential regulatory overlap between these systems (Table 1).

We tested compounds characteristic of specific infections sites, such as pH and seminal plasma, and some more generally applicable to infection, such as reactive oxygen species (ROS), to gain insight into T4SS expression in varying contexts. pH varies greatly between infection niches: engulfed bacteria encounter variable acidification by host cell phagosomes, whereas the average pH of semen is ∼8.2 (Haugen and Grotmol 1998; Lee et al. 2003). Liquid β-galactosidase assays of cultures exposed to decreased or increased media pH conditions at which *N. gonorrhoeae* can survive *in vitro* (6.3 and 8.0, respectively) showed no significant changes in TraD expression (Supplemental Figure S1).

Spermidine, an abundant polyamine in the male urogenital tract that helps gonococci persist against innate immune factors (Goytia and Shafer 2010), had no effect on TraD expression. However, whole seminal plasma as would be encountered in semen-mediated transmission of gonorrhea visibly increased TraD reporter expression on plates (Table 1).

During host infection, *N. gonorrhoeae* must combat reactive oxygen and nitrogen species produced by both the host immune response and bacterial metabolism (Archibald and Duong 1986; Cardinale and Clark 2005; Criss et al. 2009) (Reviewed in Seib et al. 2006; Winterbourn et al. 2016). TraD expression increased with proximity to a disk of 3% hydrogen peroxide (Table 1). None of the nitrogen species tested elicited a change in TraD expression (Table 1). Based on these initial experiments, we proceeded to further investigate compounds with the most profound effects on TraD, beginning with the metals iron, zinc, copper, and manganese.

### Iron availability and T4SS expression

Like many bacteria, gonococci employ a ferric uptake regulator, Fur, to control gene expression in response to intracellular iron (Berish et al. 1993). We found that a *fur* point mutant, *fur-1*, wherein iron response is compromised due to impaired functionality of Fur, had increased levels of TraD in the disk diffusion assay compared to the wild type (Table 1). Canonically, the presence of iron allows Fur monomers to dimerize, and these dimers bind promoters to repress transcription (Bagg and Neilands 1987; De Lorenzo et al. 1987).

To confirm and quantify the results from the disk diffusion assays, we performed β-galactosidase assays on *N. gonorrrhoeae* expressing *traD::lacZ*, grown in liquid cultures. These assays demonstrated that the *fur-1* strain (AKK548) exhibited a significant increase in TraD expression, with nearly 2-fold more β-galactosidase made in the *fur-1* strain than in the wild type (Figure 2A). The *fur-1* mutant showed no difference between iron replete (GCBL + 1.2 μM Fe(NO_3_)_3_) and iron deplete (GCBL + 100 μM desferal) media on β-galactosidase activity, behaving as expected for this iron-unresponsive mutant. 100 μM desferal is in excess of the amount necessary to chelate all Fe^3+^ in cGCBL and was sufficient to lead to strong β-galactosidase activity in a control strain carrying a *tbpB*::*lacZ* transcriptional fusion (data not shown). Western blots detecting an epitope-tagged TraD also showed increased TraD protein in the *fur-1* strain (Figure 2B). Transcript levels of *traD* (located on GGI operon 1) and *traK* (located on operon 2) (Figure 1A) in the *fur-1* mutant exhibited modest increases over wild type regardless of iron availability (Figure 2C), offering some explanation for the observed increase in protein. We were able to quantify secreted DNA for both the wild type and *fur-1* strain, although DNA quantification is often highly variable (Ramsey et al. 2015). The *fur-1* strain exhibited a non-significant trend of increased DNA secretion compared to the wild type (Figure 2D), indicating a potential change in T4SS activity in response to the increased protein expression.

**Figure 2.**
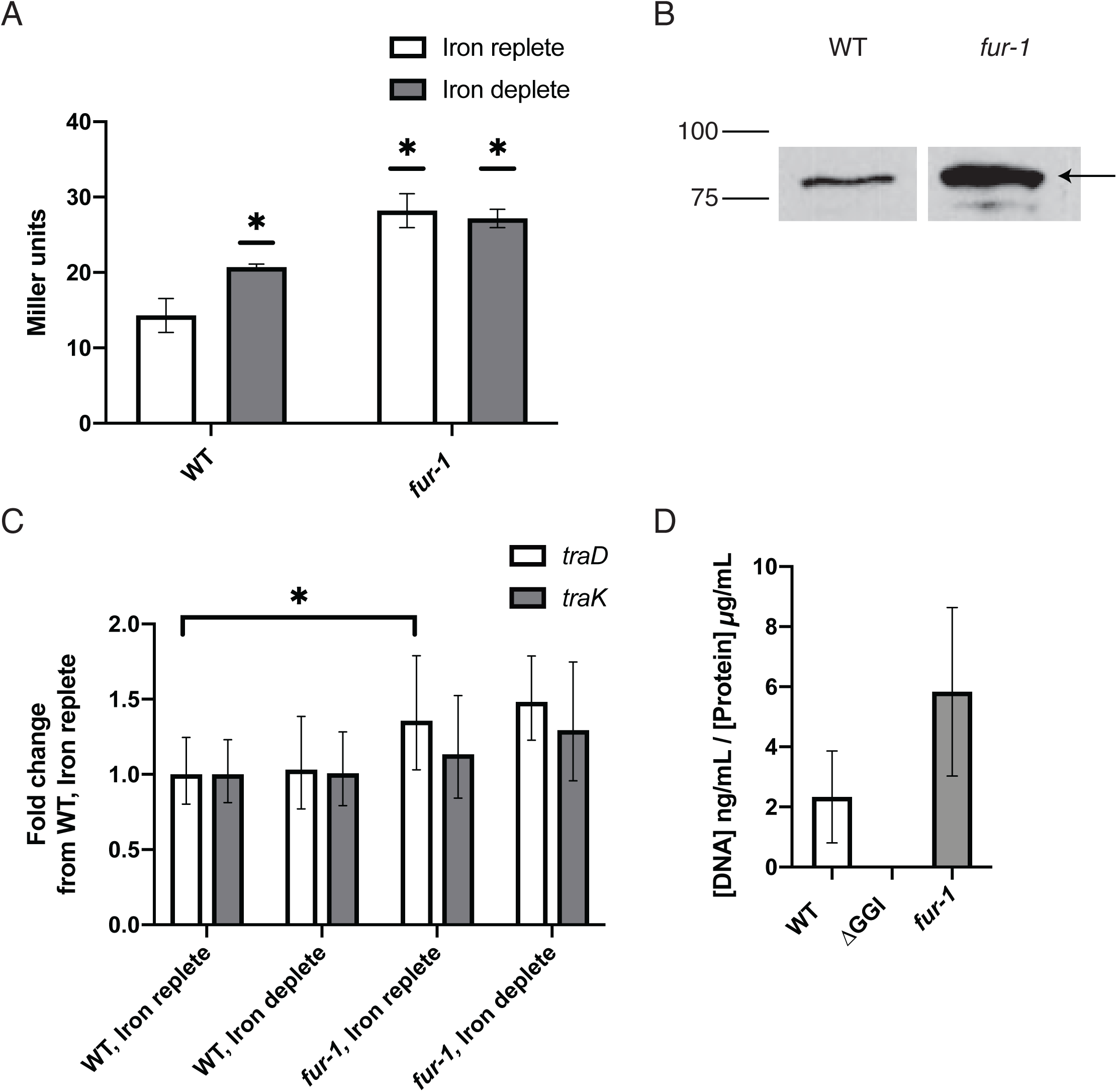
Iron sensing plays a regulatory role in T4S. (A) β-galactosidase assay detecting TraD-LacZ expression from strains wild type for *fur* vs. the *fur-1* mutant in both iron replete and iron deplete growth medium. * p<0.05 in student’s t-test comparing sample to wild type *fur* in iron replete media. Average of three experiments, error bars are SEM. (B) Representative western blot against the FLAG tag in TraD-FLAG3 strains with wild type *fur* and *fur-1* in iron replete media. Arrow indicates TraD-FLAG3 band. (C) qRT-PCR comparing GGI transcripts from wild type and the *fur-1* mutant in both iron replete (cGCBL + 1.2 μM Fe(NO_3_)_3_) and iron deplete (GCBL + 100 μM desferal) growth medium. * p<0.05 in student’s t-test comparing ΔC_T_ values to those of wild type in iron replete media. Average of three experiments, error bars are 95% confidence intervals. Transcripts normalized to *rmp.* (D) Quantification of DNA in supernatants from wild type (MS11), ΔGGI (ND500), and the *fur-1* mutant. Values corrected for extracellular DNA resultant from lysis by subtracting ΔGGI (ND500) values in each experimental replicate. Students t-test determined no significant differences compared to MS11. Average of three experiments, error bars are SEM.

Iron depletion alone only partially recapitulated expression differences observed in the *fur-1* mutant. β-galactosidase assays using the *traD::lacZ* translational reporter revealed that TraD protein was significantly more abundant under iron chelated conditions as compared to the supplemental iron condition (Figure 2A). However, for both *traD* and *traK,* transcript levels were the same in iron replete and deplete conditions (Figure 2C). These data suggest a model in which Fur plays two roles: firstly, Fur represses *traD* transcription regardless of iron availability, perhaps from low-affinity or transient promoter binding (Yu and Genco 2012); secondly, Fur and iron levels interact, perhaps through a second factor, to keep TraD protein levels low under iron replete conditions but allow TraD levels to rise during iron depletion.

### Copper, zinc, and cooperative upregulation of TraD expression

Niches occupied for gonococci vary highly in metal availability and metal concentrations are often not well known. For example, Goullé et al. found a median zinc concentration in human plasma of ∼11.1mM, whereas Owen & Katz did an extensive scan of reported literature values for zinc concentration in human semen and found it to be an order of magnitude higher, at 224 ± 53mM (Goullé et al. 2005; Owen and Katz 2005). We tested multiple concentrations of both zinc and copper, striving to add enough to elicit an effect without disrupting *in vitro* growth.

Disk diffusion assays with both ZnSO_4_ and CuSO_4_ demonstrated an increase in β-galactosidase activity for the *traD::lacZ* strain with increasing concentration of these metals. Quantitative β-galactosidase assays confirmed that supplementary zinc and copper both increased TraD production (Figure 3A and B). TraD upregulation was concentration-dependent, with the highest TraD expression observed at 250µM ZnSO_4_ and 500µM CuSO_4_ (Figure 3A and B). Zinc treatment did not significantly alter transcript levels for *traD* or *traK* (Figure 3C).

**Figure 3.**
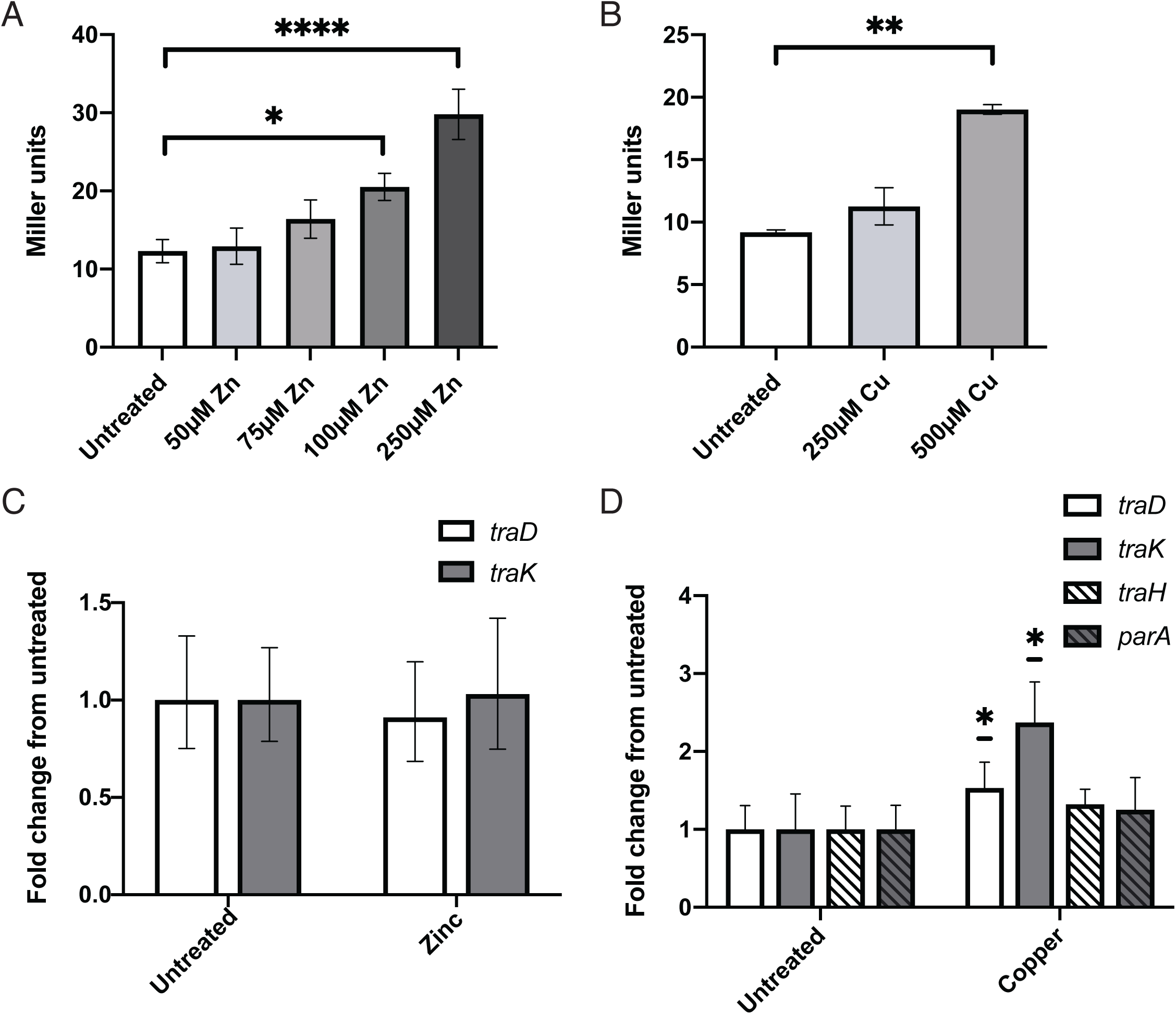
Zinc and copper both alter TraD expression. (A,B) β-galactosidase assays of *traD::lacZ* gonococci with varying concentrations of (A) ZnSO_4_ or (B) CuSO_4_. Significance determined by one-way ANOVA and Dunnett’s posttest. Average of three experiments, error bars are SEM. (C,D) qRT-PCR of GGI transcripts in response to (C) 100μM ZnSO_4_ or (D) 500μM CuSO_4_. Significance determined by Student’s t-test comparing ΔC_T_ values. Average of three experiments, error bars are 95% confidence interval. Transcripts normalized to *rpoB.* *p<0.05, **p<0.01, ****p<0.0001.

We tested the effect of copper treatment on *traD* and *traK* transcript levels, as well as those of *traH* (operon 3) and *parA* (operon 4), representing all four GGI operons necessary for secretion (Figure 1A). All of the tested transcripts exhibited small increases with copper treatment, with *traD* and *traK* being statistically significantly higher (∼1.5 and 2.3-fold, respectively) (Figure 3D). The effects of increased concentrations of zinc and copper look very similar to those of iron chelation – small (if any) transcriptional changes and more pronounced protein increases, suggesting largely post-transcriptional upregulation of TraD protein expression.

Copper and zinc toxicity have both been identified as mechanisms of bacterial killing in the phagosome (Wagner et al. 2005; White et al. 2009; Neyrolles et al. 2015; Sheldon and Skaar 2019). We next asked if zinc and copper could act cooperatively to further increase TraD expression. Since the phagosome also employs iron sequestration to kill bacteria, we included desferal in β-galactosidase assays of *traD::lacZ* with combination treatments. All binary combinations of desferal, zinc, and copper, as well as treatment with all three, significantly increased TraD expression (Figure 4A). No significant changes in growth were observed in response to these treatments, as assessed by quantifying protein concentration in the cultures after growth (data not shown). We used western blotting against the FLAG epitope tag to further explore the specificity of post-transcriptional metal regulation. Gonococci with 3XFLAG-tagged *traD* and *traK* were grown in GCBL and treated with either desferal and zinc or desferal, zinc, and copper. The *traK* construct was overexpressed from a complementation locus to provide ample transcript and allow visualization by western blot. TraD protein was distinctly increased by both metal treatments, whereas TraK had no change in abundance (Figure 4B). As with the individual metal conditions, combination desferal, zinc, and copper treatment had no significant effect of GGI transcript levels (Figure 4C).

**Figure 4.**
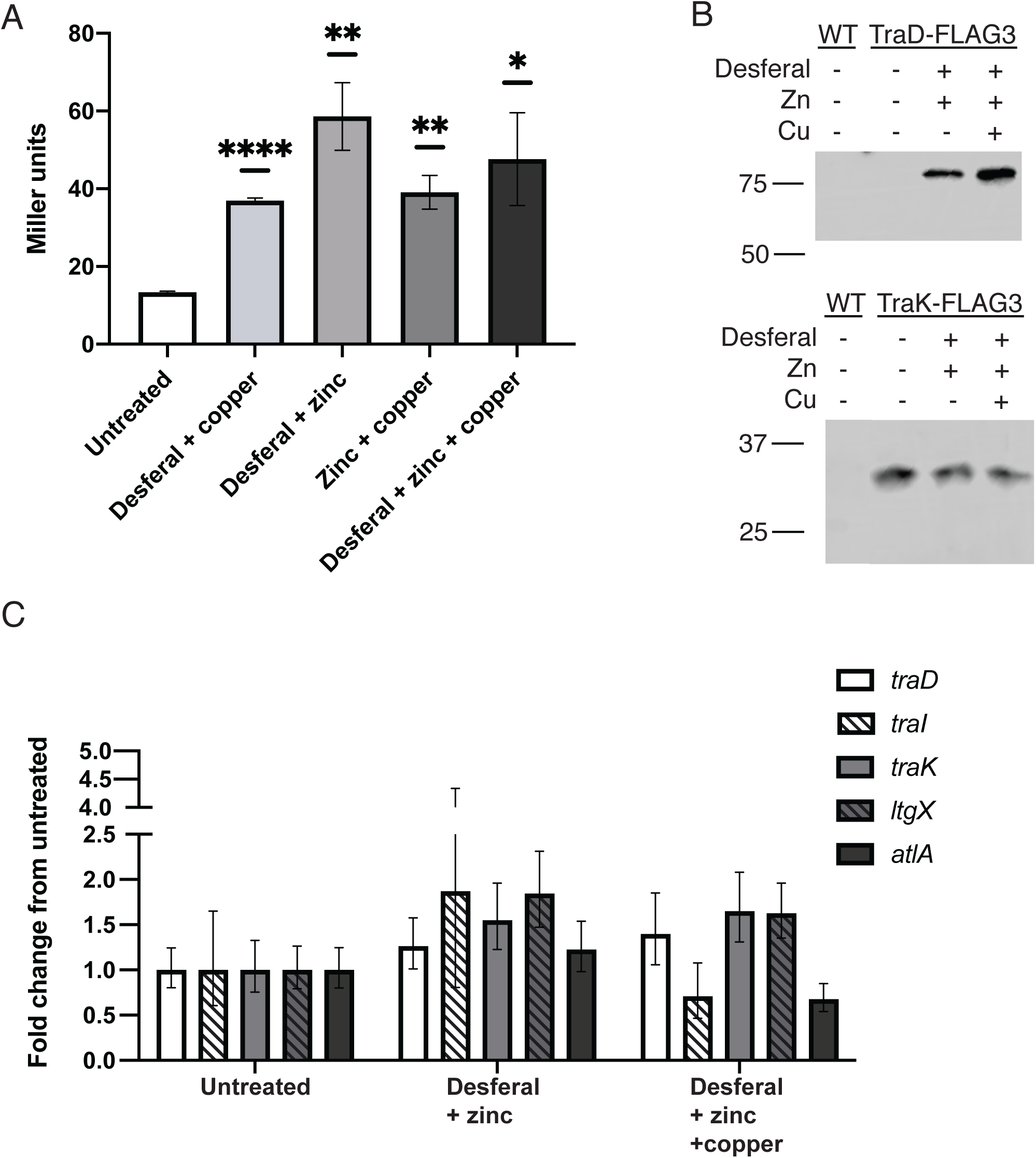
Semi-cooperative upregulation of TraD by zinc, copper, and iron chelation. 100μM desferal, 250μM ZnSO_4_, and 500μM CuSO_4_. (A) β-galactosidase assay detecting TraD-LacZ expression. Significance determined by student’s t-test comparing samples to untreated cultures. * p<0.05, **p<0.01, ****p<0.0001. Average of three experiments, error bars are SEM. (B) Representative western blot against the FLAG epitope. Native expression of *traD::FLAG3* (top), induced overexpression of *traK::FLAG3* (bottom). Wild type (WT) has no FLAG-tag. (C) qRT-PCR finds no significant differences between all tested GGI transcripts by student’s t-test comparing ΔC_T_ values. Average of four experiments, bars are 95% confidence interval. Transcripts normalized to *rpoB*.

The mechanism of toxicity for copper and zinc is often mismetallation of proteins (Sheldon and Skaar 2019; Baksh and Zamble 2020). Since Fur is an active regulator in this system, we performed β-galactosidase assays to determine whether the effects of zinc and copper on TraD expression were a result of Fur mismetallation. The *traD::lacZ fur-1* mutant strain responded to both zinc and copper treatments by expressing more TraD, indicating the these two metals do not act by disrupting the activity of Fur (Supplemental Figure S2).

### Manganese represses GGI expression

The only compound in our disk diffusion screen that downregulated TraD expression was manganese chloride (Table 1, Figure 1C). Similar to all tested metals in this study, manganese concentrations vary highly between body sites and have not been characterized in the context of *Neisseria* infections. One study reports median manganese concentrations in whole blood to be ∼138µM, while isolated plasma had only ∼20µM (Goullé et al. 2005). We performed western blotting with a TraD-FLAG3 strain (AKK556) and detected less TraD in cultures grown with supplemental manganese (cGCBL + 100µM MnCl_2_), confirming that TraD is expressed less in response to this manganese treatment (Figure 5A). Quantification of extracellular DNA in wild type cells (strain MS11) revealed that cultures treated with manganese had consistently less DNA in the supernatant than untreated cultures (Figure 5B). A GGI deletion strain (ND500) was used as a control for lysis in these experiments. Surprisingly, qRT-PCR demonstrated that transcript levels for *traD* and *traK* were significantly decreased in cultures grown with 100 μM MnCl_2_ (Figure 5C). This result indicates that manganese is working in a unique manner from the other metals identified in our screen, acting at the transcriptional level to downregulate GGI expression.

**Figure 5.**
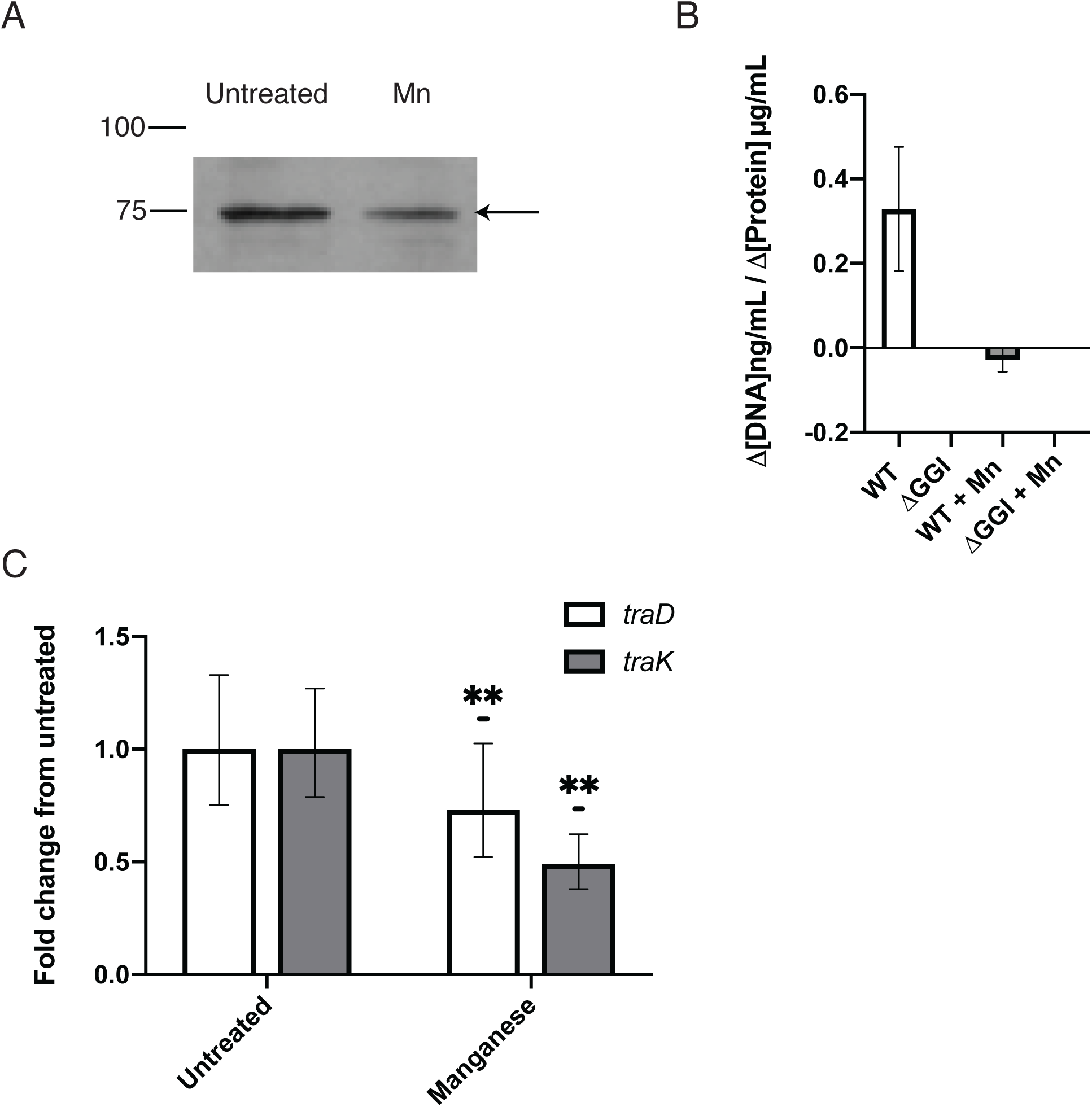
Manganese represses T4S. (A) Representative western blot for TraD-FLAG3 demonstrates a visible decrease in TraD in cultures treated with 100 μM MnCl_2_. Arrow indicates TraD-FLAG3 band. (B) Extracellular DNA quantification shows less DNA is secreted in manganese-treated cultures. Values corrected for extracellular DNA resultant from lysis by subtracting ΔGGI (ND500) values in each experimental replicate. Average of three experiments, error bars are SEM. (C) qRT-PCR shows decreased *traD* and *traK* transcript with 100 μM MnCl_2_. **p<0.01 in student’s t-test comparing ΔC_T_ values. Average of three experiments, error bars are 95% confidence interval. Transcripts normalized to *rpoB*.

### GGI genes are expressed during urethral infection

Many of the factors that we have found to alter T4SS protein expression are physiologically relevant to gonorrhea infection. We postulate that the low expression levels observed under standard laboratory conditions are not representative of expression in the human host. Furthermore, most of the effects that were identified in the compound screen appear to act at the post-transcriptional level. This finding was somewhat surprising and made us question whether there are noteworthy changes in GGI transcripts between the lab and the host environment. A recent study from Nudel *et al*. collected urethral swabs from infected males and performed RNA-Seq on both the *N. gonorrhoeae* isolate during host infection and as a monoculture in laboratory chemically-defined medium CDM (GEO accession GSE113290) (Nudel et al. 2018). We identified five GGI^+^ urethral isolates in the resultant dataset (Supplemental Table S3) and used Rockhopper (Mcclure et al. 2013; Tjaden 2015) to compare GGI expression in the urethral swabs and growth of the pure isolates in medium.

The GGI exhibited distinct gene expression profiles between laboratory culture and urethral infection (Figure 6). Of the 62 GGI genes detected by Rockhopper, 35 (∼56%) had significantly higher expression during urethral infection. Furthermore, 20 of the 21 genes that are essential for T4SS-dependent DNA secretion were more highly expressed in the urethral infections, half of these significantly so. The exception, *traH*, had significantly higher expression in culture medium though was still expressed at detectable levels during infection.

**Figure 6.**
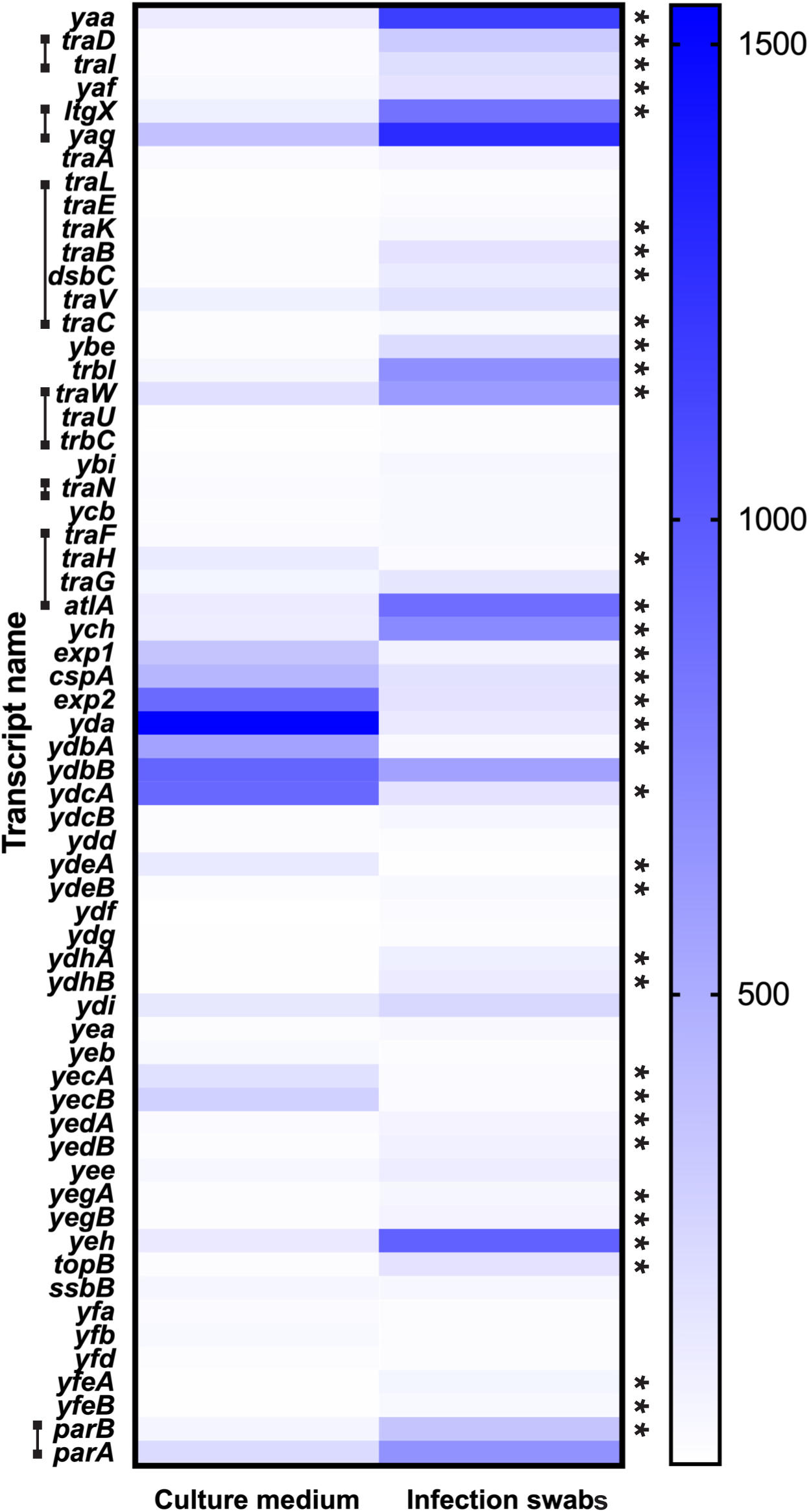
The GGI is differentially expressed during natural urethral infection in humans. Heat map of GGI gene expression in laboratory grown gonococcal isolates and urethral swab samples. Brackets on the left indicate genes necessary for DNA secretion in laboratory culture. n=5. *q-value<0.05.

The portion of the GGI spanning *ych-ydcA*, which consists largely of genes of unknown function, has been previously demonstrated to be dispensable for DNA secretion (Pachulec et al. 2014). This region, spanning ∼4.4 kb, was notably downregulated during urethral infection compared to laboratory growth conditions (Figure 6). Furthermore, Rockhopper identified several novel antisense transcripts in this region, and many of these antisense transcripts were significantly increased in the urethral swab samples. Hypothesizing that this region might contain genes that negatively regulate the T4SS, we deleted the *exp1*-*ydcA* region from the gonococcal chromosome. However, measurements of *traK* and *parA* expression did not show increased transcript levels in the mutant. We conclude from the RNA-Seq analysis that while the T4SS is not robustly expressed in the laboratory, its transcription is upregulated during infection of the male urethra in a specific and consistent manner.

### Surface adherence enhances GGI expression

At some infection sites gonococci are capable of biofilm formation, creating a matrix consisting partly of extracellular DNA atop epithelial cells (Greiner et al. 2005; Steichen et al. 2008; Steichen et al. 2011). The gonococcal T4SS has been shown to contribute to biofilm establishment, although it is not essential for biofilm formation (Zweig et al. 2014). However, the effect of the biofilm environment on T4SS expression has not heretofore been investigated. We grew the bacteria in stationary 12-well tissue culture plates for 6, 12, and 24 hours to elucidate potential effects of planktonic vs. surface-adhered cellular lifestyles on the secretion system. Planktonic cells were separated from cells adhered to the bottom of the wells by gentle pipetting, and then the two populations of cells were analyzed in parallel. Due to the limited number of cells obtained from growth in this manner, we were unable to detect β-galactosidase activity from the *traD::lacZ* reporter strain in either planktonic or surface-adhered cells. However, qRT-PCR indicated that adhered cells had significantly increased transcript levels at 6 and 12 hours of stationary growth, with at least a 4-fold increase in transcript level over their planktonic counterparts (Figure 7). Increases were less pronounced, but still over 2-fold, after 24hours of growth. Cellular lifestyle thus appears to exhibit a regulatory effect on GGI transcription, with growth in adherent conditions increasing T4SS expression especially at early time points. This observation is relevant to infection at many host sites, where gonococci adhere to host cells and form biofilms on host tissues.

**Figure 7.**
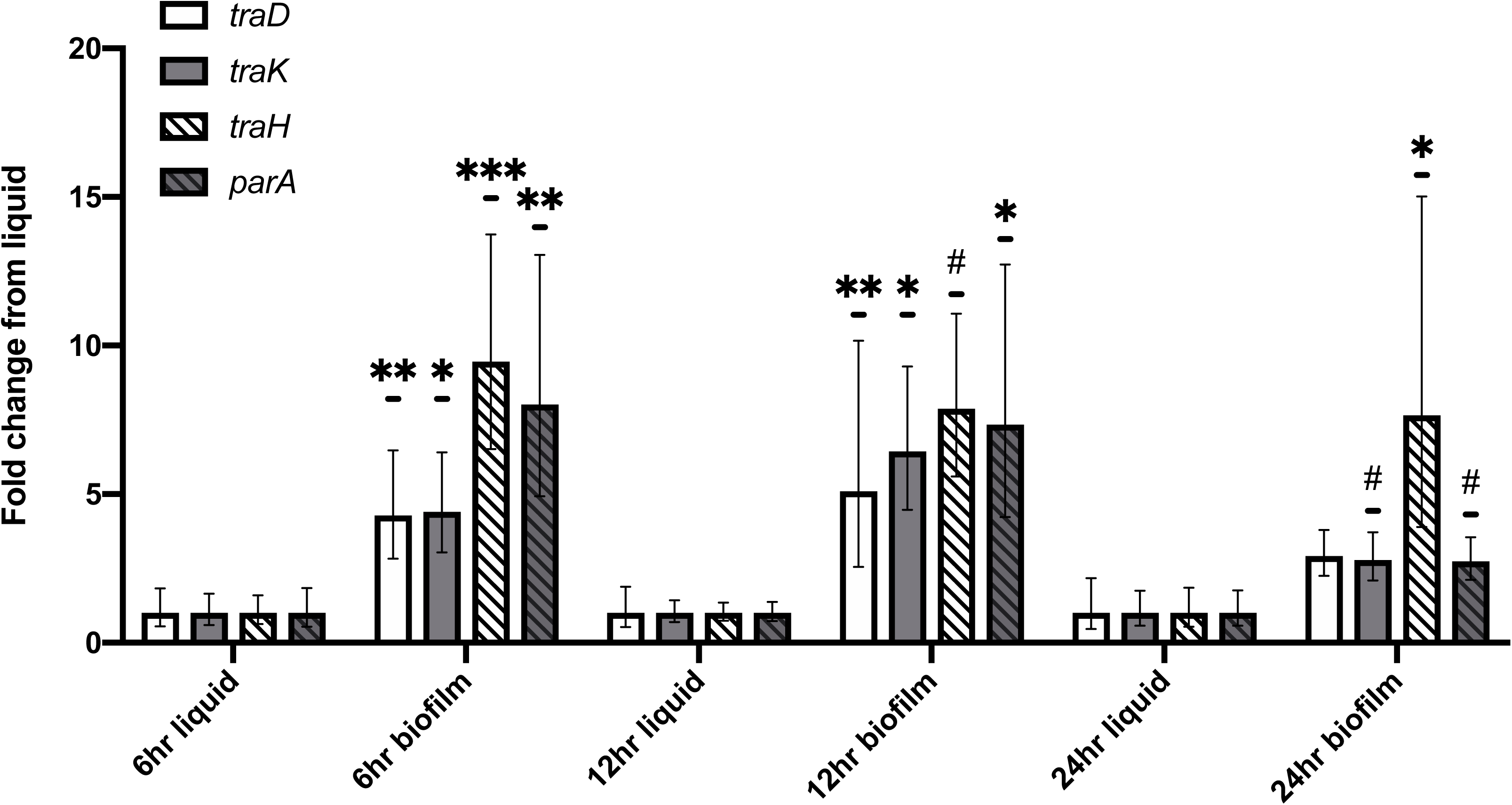
GGI transcript levels are elevated in surface-adhered cells. qRT-PCR of representative GGI genes. # p<0.1, * p<0.05, **p<0.01, ***p<0.001 in student’s t-test comparing ΔC_T_ values. Average of four experiments, error bars are 95% confidence interval. Transcripts normalized to *rpoB*.

## Discussion

Type IV secretion systems are a diverse family of protein machines that play myriad roles in host-pathogen interactions. The effects of the gonococcal T4SS have been observed both in the laboratory and at the population level, however observable expression of this system has been low (Hamilton et al. 2001; Hamilton et al. 2005; Ramsey et al. 2014; Harrison et al. 2016; Shockey 2019). We expect that the gonococcal T4SS might be assembled or otherwise activated by specific environmental signals during infection, especially because it is likely a costly apparatus to build and run. In this study, we investigated gonococcal responses to conditions that might be encountered during infection and bacterial regulatory machinery that alter expression or activity of the T4SS.

### Upregulation of T4SS protein expression by phagosome-like metal conditions

After testing a variety of compounds, we found effects on GGI protein expression from four metals – manganese, iron, zinc, and copper – some acting at the transcriptional and others at the post-transcriptional level. Manganese depressed TraD expression at the transcriptional level, and repressed *traK* transcript levels (operon 2) as well.

Iron sensing and response in gonococci is controlled by the ferric uptake regulator, Fur. Fur dimerizes when iron is bound and binds Fur-boxes at promoters, where it can to directly repress transcription, directly activate transcription, or indirectly activate transcription by repressing a repressor (Yu et al. 2016). The *fur-1* gonococcal strain, which has a point mutation in the *fur* gene leading to partially functional Fur (Thomas and Sparling 1996) had significantly increased TraD protein. Transcript levels for *traD* were modestly increased in the *fur-1* mutant compared to wild type *N. gonorrhoeae* under iron replete conditions (Fig 2C), suggesting Fur slightly represses *traD* transcription. However, iron chelation by desferal treatment had no effect on *traD* transcription. These data may be explained by transient binding of Fur monomers to AT-rich promoters in the GGI. Apo-Fur repression, in which Fur acts independent of iron, has been shown to play a role in the Fur regulon of gonococci, and the Fur box sequence is variable and AT-rich (Yu and Genco 2012; McClure et al. 2020) (Figures 2C). Interestingly, iron deplete cultures did have significantly increased TraD protein expression in the LacZ reporter assay. Increased levels of TraD::FLAG3 were also detected in response to iron chelation.

Synthesizing these data, we suggest a model (Figure 8) in which iron-dependent Fur regulation acts at the post-transcriptional or translational level, but does not affect *traD* transcription. This could be achieved through an intermediate regulator such as a protein or small RNA (sRNA). These regulatory elements have been identified as mechanisms of Fur activity previously in the pathogenic *Neisseria* (Mellin et al. 2007; Yu et al. 2016; Jackson et al. 2017).

**Figure 8.**
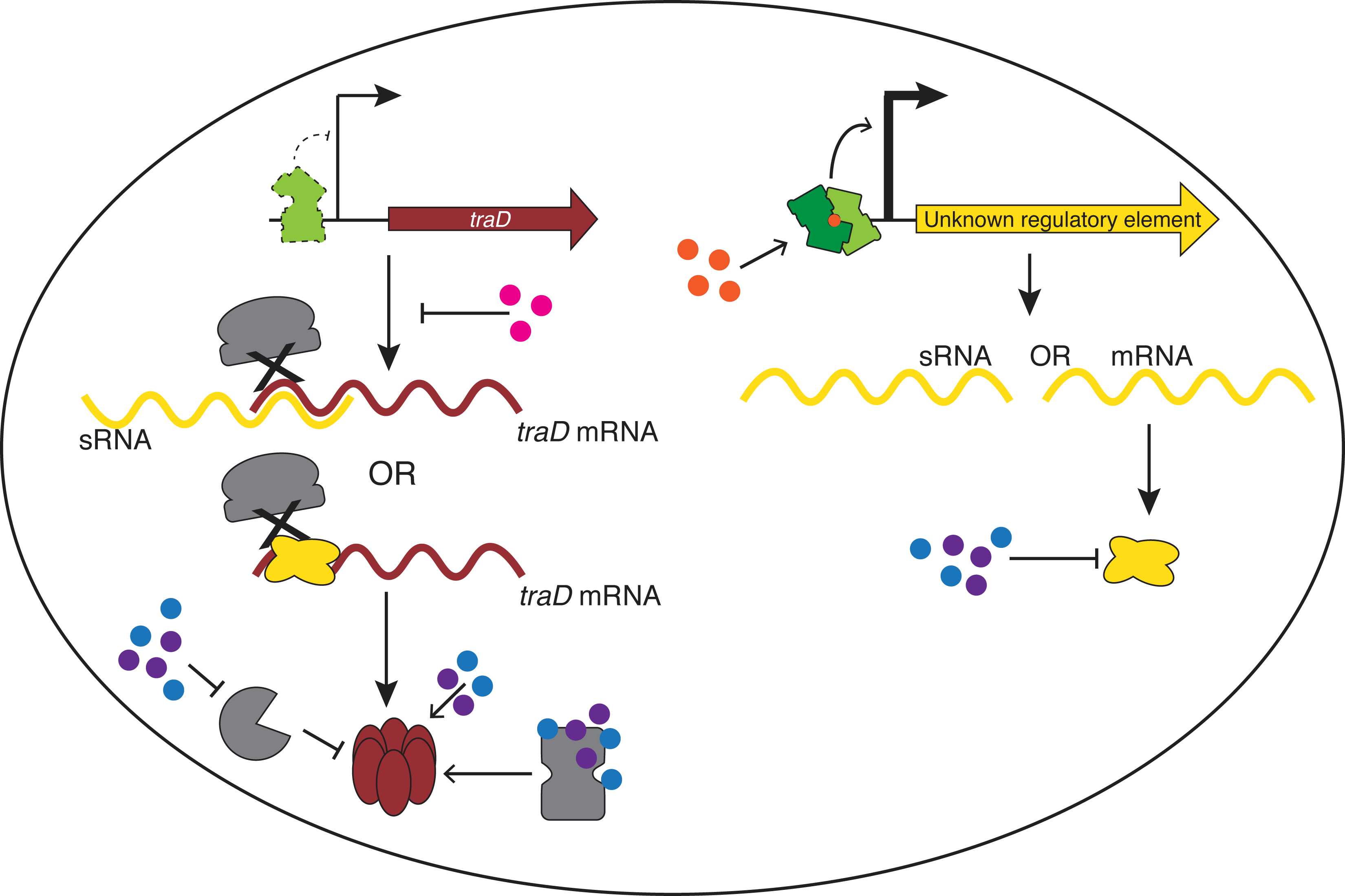
Model of TraD regulation by metals. When iron (orange) availability is high, Fur represses expression of the type IV secretion system coupling protein, TraD, through an unknown intermediate (yellow) (depicted here as either an sRNA or protein). In the phagosome, iron sequestration releases Fur-mediated TraD repression, resulting in increased TraD protein levels. Residual apo-Fur binding may happen transiently at the *traD* promoter, with small effects on *traD* transcript levels. High levels of manganese (pink) repress *traD* transcription, but the phagosome pumps manganese away from intracellular gonococci allowing transcription to proceed. Zinc (purple) and copper (blue) contribute to the upregulation of TraD in a undefined manner; here we depict several points at which this regulation could occur: repression of a repressor or protease, direct stabilization of TraD, or enhancers of a TraD chaperone.

Apart from regulation by manganese and iron, both zinc and copper supplementation increased TraD protein levels. Zinc elicited no transcriptional changes, whereas copper yielded small but significant increases in *traD* and *traK*, suggesting that these metals act differently within the GGI regulon (Fig 3C and D). Pairwise combinations of desferal, zinc, and copper treatments, as well as treatment with all three, all enhanced TraD expression beyond that of individual treatments (Fig 4A). Zinc and copper could be acting at a number of points in the regulatory network: they could enhance TraD protein stability, perhaps by inhibiting the activity of a protease or by activating a chaperone protein for TraD, or they could act upon another regulator within this system, potentially one that is also controlled by iron-dependent Fur regulation. (Baksh and Zamble 2020) (Figure 8).

Our finding that TraD protein expression is increased specifically and semi-cooperatively by iron chelation combined with zinc and copper indicate that there may exist a specific and complex infection niche in which gonococci activate the T4SS in response to these combined signals. The host environment during symptomatic gonorrhea infection is highly inflammatory;host response to gonococci usually includes an influx of macrophages and neutrophils, in which gonococci are capable of surviving after engulfment (King et al. 1978; Casey et al. 1986; Apicella et al. 1996; Criss et al. 2009; Reviewed in Palmer and Criss 2018). In both types of immune cell, the phagosome contains active mechanisms such as NRAMP1 to pump iron and manganese out of the phagosome, using nutritional immunity strategies to kill bacteria (Jabado et al. 2000; Canonne-Hergaux et al. 2002). A transcriptional response to facilitate iron acquisition has been recorded in response to professional phagocytes and growth during human infection (Zughaier et al. 2014; McClure et al. 2015). At the same time, the toxic cations zinc and copper (observed only in macrophages to date) are pumped into the phagosome (Wagner et al. 2005; Botella et al. 2011; Neyrolles et al. 2015; Ong et al. 2018; Sheldon and Skaar 2019). Sensing and responding to these types of stressors is essential for maintaining metal homeostasis, and a variety of mechanisms and responses have been observed in bacteria (Chandrangsu et al. 2017). We propose that upon engulfment by a host immune cell, *N. gonorrhoeae* in the phagosome respond to these specific metal conditions by synthesizing TraD, and potentially other T4SS proteins. As TraD is the coupling protein, it could then recruit substrates for secretion by the T4SS. Protein stoichiometry has not been elucidated in this system; it is possible that TraD is a limiting factor for T4S that is turned on upon entrance into the host cell phagosome for immediate secretion of DNA or possibly protein effectors.

Expressing the T4SS at this time could confer several advantages to the bacterial invader. Firstly, every new host has the possibility of harboring new bacterial populations, both existing populations of *N. gonorrhoeae* (especially in cases of asymptomatic gonorrhea) and commensal microflora. These present an opportunity for T4SS-mediated horizontal gene transfer, which would enhance survival of gonococcal genes, including GGI genes, beyond the duration of gonococcal infection. Additionally, one could imagine that in the face of toxic cationic metals and host defense molecules, a bacterium surrounded by secreted, negatively charged DNA may be less susceptible to killing.

The gonococcal T4SS was previously noted to be expressed during intracellular infection of epithelial cells in tissue culture. Zola et al. found that gonococcal mutants lacking *tonB* could survive if they carried the GGI, whereas strains lacking the GGI would not survive intracellularly (Zola et al. 2010). Furthermore, mutations affecting structural components of the T4SS eliminated intracellular survival in the absence of TonB. Since all gonococci in the wild do express TonB and use it for metal acquisition, the significance of the ability of GGI^+^ gonococci to bypass TonB was not clear. However, the current findings suggesting that phagosome-like metal conditions upregulate type IV secretion, together with the knowledge that the T4SS is expressed during intracellular growth, suggest that the phagosome may be a previously unappreciated niche for gonococcal type IV secretion. VirB-type T4SSs have been noted to play important roles in intracellular growth for multiple pathogens such as *Legionella pneumophila* and *Brucella abortus*, and a T4SS inner membrane complex in nontypeable *Haemophilus influenzae* facilitates the release of DNA and proteins needed to form biofilms (Berger and Isberg 1993; Vogel et al. 1998; O’Callaghan et al. 1999; Celli et al. 2003; Jurcisek et al. 2017).

### Increased GGI expression during urethral infection and surface adherence

We investigated transcriptional changes during urethral infection and found that the transcriptional profile of the GGI was markedly increased in urethral infection isolates of *N. gonorrhoeae* compared to those same isolates grown in laboratory media (Figure 6, GEO accession number GSE113290). The upregulation of almost all genes essential for DNA secretion indicate an increase in T4SS expression during urethral infection. The increased T4SS gene expression detected in urethral swabs could be representative of intra- or extracellular gonococci, or both. Extracellular gonococci often live in biofilms, observed on the cervix during naturally occurring infections (Greiner et al. 2005; Steichen et al. 2008). Based upon the significantly higher GGI transcript levels we found in newly surface-adhered cells and the transcriptomic profiles of urethral infection swabs, we suggest that adherence triggers GGI gene expression and allows for DNA secretion. Secreted DNA has been shown to nucleate biofilm formation, and within biofilms gonococci have higher rates of horizontal gene transfer (Zweig et al. 2014; Kouzel et al. 2015). Upon immune cell phagocytosis, the combined metal sequestration and toxicity mechanisms act upon the abundant GGI transcripts to increase specific protein levels, including TraD, further upregulating T4SS expression and activity in intracellular gonococci.

This study has identified several environmental factors that work in combination to regulate the gonococcal T4SS. Regulation appears to act at multiple levels, potentially differing between intracellular and extracellular bacteria. Metal regulators map onto phagosomal conditions, which involve a combination of nutrient sequestration and metal toxicity to kill invading bacteria. Additionally, surface adherence increases GGI expression, and may be a contributing factor for the robust upregulation of the GGI observed during urethral infection. The specific and complex upregulation of the T4SS within multiple infection niches indicate that the T4SS is part of the gonococcal response to the human host and points to a potential role for this system in the infection process.

## Experimental Procedures

### Bacteria and growth conditions

Gonococci were grown on GCB agar plates with Kellogg’s supplements, in liquid GCBL medium with Kellogg’s supplements and 0.042% sodium bicarbonate (called “complete GCBL” or “cGCBL”) (Kellogg et al. 1963), or in Graver-Wade (GW) medium (Wade and Graver 2007). Bacterial strains used in this study are listed in Supplementary Table S1. All mutants are derived from the wild type strain MS11. Primer sequences are listed in Supplementary Table S2. Where needed, kanamycin was used at 40 μg mL^-1^ for *E. coli* and 80 μg mL^-1^ for *N. gonorrhoeae.* Unless otherwise noted, desferal was used at 100 μM and metals were used at the following concentrations: 100 μM MnCl_2_, 250 μM ZnSO_4_, 500 μM CuSO_4_.

### Construction of traD::lacZ reporter strain

The 3’ region of *traD* was PCR-amplified from MS11 chromosomal DNA with primers PstI-traDlacZ-F and 76-R, then digested with *Hin*dIII and *Pst*I, resulting in a 0.75kb fragment. This fragment was ligated into the *Hin*dIII and *Pst*I sites of pPK1008, generating pKL5 which contained a ‘*traD-lacZ* translational fusion. DNA downstream from *traD*, containing *yaa* truncated at the 5’ end, was PCR-amplified with primers PspOMI-83F and 86-R, digested with *Psp*OMI and *Spe*I and ligated into the *Not*I and *Spe*I sites in pKL5, generating pKL10. The *aph3* kanamycin resistance gene from pKH99 was cloned into the *Spe*I and *Age*I sites in pKL10, immediately downstream from the *traD-lacZ* fusion, generating the reporter plasmid pAKK1. *N. gonorrhoeae* strain MS11 was transformed with linearized pAKK1, and transformants containing the *traD-lacZ* reporter recombined in the GGI were selected on GCB agar containing Kan, resulting in strain AKK500. The *aph3* gene was removed by transforming AKK500 with pKL10 and screening for transformants that were sensitive to Kan, resulting in reporter strain AKK520.

### Construction of a TraD-FLAG3 N. gonorrhoeae strain

To create the *traD-FLAG3* construct, a 0.27kb fragment containing a linker fused to the triple epitope FLAG tag was obtained from pMR100 by cutting with *Pst*I, blunting with T4 DNA polymerase, then digesting with *Xba*I. This fragment was cloned into the *Sma*I/*Spe*I sites of pAKK1, resulting in the final C-terminal FLAG3-tagged TraD construct (pAKK28). *N. gonorrhoeae* MS11 was transformed with pAKK28 resulting in AKK556 with TraD-FLAG3 at the native *traD* site.

### Construction of fur-1 mutants

The *fur-1* allele from *N. gonorrhoeae* MCV403 (gift from C. Cornelissen) was PCR-amplified with primers fur_up-F and fur_down-R and transformed into *N. gonorrhoeae* MS11, AKK520, and AKK556. Putative transformants were identified by their smaller colony size and confirmed by sequencing the *fur* gene. The *fur-1* allele in MCV403 contains a tyrosine 82 to cysteine substitution, which was present in the original *fur-1* mutant isolated in (Thomas and Sparling 1996). Our sequencing of the *fur* allele in MCV403 revealed it had two additional mutations: a silent ATT to ATC base change within isoleucine 7 and a GAG to GGG mutation changing glutamate 48 to glycine. The *fur-1* alleles in AKK550, AKK548 (*traD::lacZ*) and AKK573 (*traD-FLAG3*) contain all three mutations.

### Disk diffusion and blue/white screening

AKK520 was grown overnight from frozen stocks on GCB agar plates, 16-20 hours. Bacteria were swabbed into 2-4 mL cGCBL and serially diluted to 10^-5^. 80 µL of the 10^-4^ and/or 10^-5^ dilutions were plated on GCB + 40 µg mL^-1^ Xgal plates. A 0.25-inch disk (Hardy Diagnostics) was placed near the bottom of the plate and suffused with 10 µL of the test compound. Plates were incubated at 37°C with 5% CO_2_. Blue color was assessed after 2 days.

### RNA-Seq data analysis

We obtained the publicly available PacBio whole genome sequence data from (Nudel et al. 2018) (BioProject PRJNA329501, Gene Expression Omnibus accession number GSE113290). We assembled the raw sequence data using Flye 2.7 with two rounds of polishing (Kolmogorov et al. 2019). We evaluated assembly quality using QUAST (Mikheenko et al. 2018). Using BLAST 2.9.0+ (Altschul et al. 1990), we identified GGI genes in each *de novo* assembly; individual GGI gene sequences from reference strain NCCP11945 (GenBank accession number CP001050.1) were used as the query sequences (Chung et al. 2008).

We proceeded with transcriptomic analysis using the GGI^+^ isolates from male patients only, as the GGI was identified in only one cervicovaginal lavage sample. Reads were aligned to the genome of NCCP11945 using Rockhopper with default parameters (Mcclure et al. 2013; Tjaden 2015). Gene expression is reported as RPKM. GGI genes with a q-value ≤0.05 were considered significantly different between groups.

### β-galactosidase assays

These assays were performed as described previously in (Ramsey et al. 2015). Cultures were started from 16-20 hours of *N. gonorrhoeae* overnight growth on GCB plates suspended in cGCBL at an OD_540_ ∼0.25 and grown for 3 hours with aeration with treatments as noted in the text. For assays testing change in media pH, gonococci were grown as above in 3 mL cGCBL pH 7.3 for 2 hours. Bacteria were collected by centrifugation and resuspended in 3 mL of fresh cGCBL at pH 6.3, 7.3, or 8.0, then grown 1 hour. After growth, cultures were placed on ice 20 minutes, and 0.5 mL was removed for protein quantification via Bradford assay. After chilling, 2 mL of culture was harvested by centrifugation, resuspended in Z buffer, and exposed to ONPG in 96 well plates. Absorbance was read with a BioTek Synergy HT plate reader. Protein concentration was used in place of optical density in the Miller equation.

### qRT-PCR

*N. gonorrhoeae* cultures in cGCBL were grown 3 hours at 37°C with rotation from an OD_540_ of ∼0.25. Where noted, static growth samples were prepared as in the static surface adherence assay. A volume of 0.75 mL of culture was combined with 0.75 mL −20°C methanol and centrifuged at 17×1000g for 1min. This procedure was repeated a total of two times, to harvest cell from 1.5 mL of culture. RNA isolation was performed as described previously (Ramsey et al. 2015). DNA was removed with the TURBO DNA-*free* kit (Invitrogen) and cDNA prepared using the iScript cDNA Synthesis kit (BioRad). qRT-PCR was performed using SYBR green reagents (BioRad) and the ΔΔC_T_ method (Applied Biosystems 1997). Statistical significance was determined with the two-tailed Student’s t-test comparing ΔC_T_ statistics according to (Yuan et al. 2006).

### Static surface adherence assays

*N. gonorrhoeae* cells from overnight growth were swabbed then suspended in cGCBL and adjusted to OD_540nm_ ∼0.1. A volume of 1 mL of culture was transferred to a sterile 12-well tissue culture plate in triplicate. The plate lid was coated in Triton solution and parafilm used to seal. The plate was incubated at 37°C + 5% CO_2_ for 6, 12, or 24 hours without agitation. To separate planktonic from adhered cells, the plate was tilted and liquid gently removed from wells with a pipette, constituting the liquid (or planktonic) fraction. A volume of 1 mL of GCBL was added to each well and adhered cells resuspended using a cell scraper and agitation with a pipetman. These cells were then collected as the surface-adhered (or biofilm) fraction.

### DNA secretion assays

These assays were performed similarly to assays described previously (Hamilton et al. 2005; Salgado-Pabón et al. 2007). Briefly, gonococci were grown on GCB agar plates overnight from frozen stocks of piliated bacteria. A piliated colony was re-streaked if reversion to non-piliation was observed on the plate. Bacteria were swabbed into GW medium, prepared without spermidine, and OD_540nm_ adjusted to 0.15-0.2. Cultures (3 mL) were grown for 1.5 hours at 37°C in 15 mL falcon tubes with rotation. After initial growth, 0.5 mL of culture was diluted into 2.5 mL of fresh GW, vortexed thoroughly, and then 0.5 mL were removed for analysis. The remaining culture was grown as before for 2 hours, at which time another 0.5 mL were removed for analysis. Samples were harvested by centrifugation for 2 minutes at 17×1000g immediately after collection. A 0.4 mL supernatant sample was removed to quantify DNA using Quanti-iT Picogreen fluorescent dye (Life Technologies). Remaining liquid and pellet were resuspended in water and protein quantified using the Bio-Rad protein assay.

### Western blots

#### Detection of TraD-FLAG3 in wild-type, manganese-treated, and *fur-1* strains

To assay wild-type vs. *fur-*1 TraD expression, overnight growth of *N. gonorrhoeae* AKK556 and AKK573 on GCB agar plates was swabbed into GCBL. To assay the effects of manganese treatment, overnight growth of AKK556 on GCB agar plates was swabbed into cGCBL, OD_540_ adjusted to ∼0.2, and 3 mL cultures were grown with rotation at 37°C for 3 hrs. For all assays, cells were collected by centrifugation and resuspended in sterile milliQ water and sonicated to obtain whole cell lysates. Protein was quantified by a Bradford assay (Bio-Rad), and equivalent quantities were electrophoresed on 10% SDS-PAGE gels. Proteins were transferred to a polyvinylidene difluoride (PVDF) membrane and blocked overnight with 5% non-fat dry milk in Tris-buffered saline containing 0.5% Tween 20 (TBST). Membranes were incubated for 1 hour with primary antibody M2 α-FLAG (Sigma) at a 1:10,000 dilution, and then washed 3 times with TBST. Blots were then incubated with goat anti-mouse peroxidase-conjugated antibody (1:10,000 dilution) (Santa Cruz Biotechnology), washed 4 times with TTBS, and then developed with the Immobilon Western chemiluminescent HRP substrate (Millipore), and exposed to film.

#### Detection of TraD-FLAG3 following desferal, copper, and zinc treatment

Liquid *N. gonorrhoeae* cultures were inoculated from 16-20 hours overnight growth on GCB agar plates into cGCBL at an OD_540_ of ∼0.25 and grown for 3 hours at 37°C with rotation. 3 mL cultures were pelleted by centrifugation in a swinging bucket tabletop centrifuge (Jouan C3i) in their entirety, washed once with cold PBS, resuspended in 500 μL water and sonicated. Lysate samples were resolved on pre-cast 10% SDS-PAGE gels (Bio-Rad). Proteins were transferred to a polyvinylidene difluoride (PDVF) membrane, blocked with 5% milk in TBST for 1 hour, and incubated with 1:20,000 M2 antibody overnight at 4°C. Secondary antibody goat α-mouse HRP were diluted 1:20,000 and administered for 1 hour. Blots were developed using the Licor Odyssey^®^ Fc imaging system.

## Acknowledgements

This work was funded by NIH grant R01AI047958. We thank Dr. Cornelissen for the *fur-1* mutant. The authors declare no conflict of interest.

## Author contributions

JPD conceived of the study. MMC, AKK, and JK acquired, analyzed, and interpreted data. AKK built strains and plasmids. ACP, aided by CSP, determined GGI presence/absence and assisted RNA-Seq analysis. MMC wrote the manuscript. JPD and AKK provided critical reading and revision.

## Statement of Data Availability

The data that support the findings of this study are available in the Gene Expression Omnibus, BioProject PRJNA329501, accession number GSE113290 at https://www.ncbi.nlm.nih.gov/geo/query/acc.cgi?acc=GSE113290. Additional data are available from the corresponding author upon reasonable request.

[dataset]Nudel K, McClure R, Genco C; (2018) Gender Specific Differences in Expression of Neisseria gonorrhoeae Antibiotic Resistance Genes During Human Mucosal Infection; Gene Expression Omnibus; GSE113290

